# Evolutionary remodeling of the ventral retina enables aerial vision in the four-eyed fish *Anableps anableps*

**DOI:** 10.1101/2025.11.02.686143

**Authors:** Louise Perez, Lina Shi, Alyssa Daspit, Keyla Pruett, Carsyn Smith, Anna Prideaux, Josane F. Sousa, Patricia N. Schneider

## Abstract

The emergence of evolutionary novelties remains a central challenge in biology, particularly for complex organs such as the vertebrate eye. While visual systems have repeatedly undergone degeneration or loss, examples of innovation are rare and poorly understood. Mechanistic insight into such gains requires tractable systems in which novel visual functions have evolved. Here we leverage the four-eyed fish *Anableps anableps* to uncover the cellular and molecular basis of adaptation to simultaneous aerial and aquatic light. Combining single-nucleus and spatial transcriptomics, we show that the *Anableps* ventral retina has been developmentally and functionally reconfigured for aerial vision. In fully aquatic teleosts such as zebrafish, ventral photoreceptors are tuned to red-shifted, downwelling light. In contrast, we show that *Anableps* exhibits an expanded and diversified ventral cone population encompassing ultraviolet-, blue-, green-, and red-sensitive types, enabling broad-spectrum, high-intensity vision from above water. To estimate *Anableps* opsin spectral properties, we employed OPTICS, a machine-learning model of opsin spectral tuning, which predicts that the ventrally expressed *opn1mw2* opsin is red-shifted relative to typical green-sensitive pigments. Analysis of dorsal-ventral patterning markers reveals an overrepresentation of ventral retinal cell types. Consistent with this, comparative spatial transcriptomic profiling of developing and adult *Anableps* and the killifish (*Fundulus heteroclitus*) indicates a dorsal displacement of the dorsal-ventral boundary in *Anableps*, resulting in a proportionally larger ventral retinal domain. Collectively, our results reveal that *Anableps* achieved its unique dual visual capacity by developmentally expanding and functionally reprogramming the ventral retina for aerial light detection, an evolutionary innovation that redefines how retinal domains can adapt to radically different visual environments.

## Introduction

While many vertebrate eye specializations involve reduction or loss of visual structures, such as seen in cavefishes^1^, species within the *Anableps* genus present a striking example of anatomical and functional innovation. The *Anableps* possesses divided pupils and corneas (Fig. 1a) and a single retina that simultaneously receives input from both air and water^2-4^ (Fig. 1b). Phylogenetically, *Anableps* belongs to a clade of Cyprinodontiform fishes, including the one-sided livebearer (*Jenynsia onca*), the killifish (*Fundulus heteroclitus*), the guppy (*Poecilia reticulata*) and the mummichog (*Fundulus heteroclitus*), all of which are fully aquatic and possess typical teleost retinas (Fig. 1c). Therefore, the unique ocular morphology of *Anableps* evolved within a lineage otherwise characterized by conventional aquatic vision, sometime the split between *Anableps* and its sister lineage *Jenynsia* to the Miocene, ∼12.5 Mya^5,6^ (Fig. 1c). In most teleosts, the retina is regionally specialized along the dorsoventral axis to match the optical properties of the aquatic environment. In species such as the zebrafish (*Danio rerio*), the ventral retina receives downwelling light filtered through the water column, which is enriched in longer wavelengths, and accordingly expresses red-shifted opsin paralogs such as *opn1mw4* and *opn1lw1*^7^. This configuration optimizes color sensitivity for the underwater visual field. In *Anableps*, however, the ventral retina faces the aerial environment, while the dorsal retina continues to process aquatic light.

**Fig. 1.**
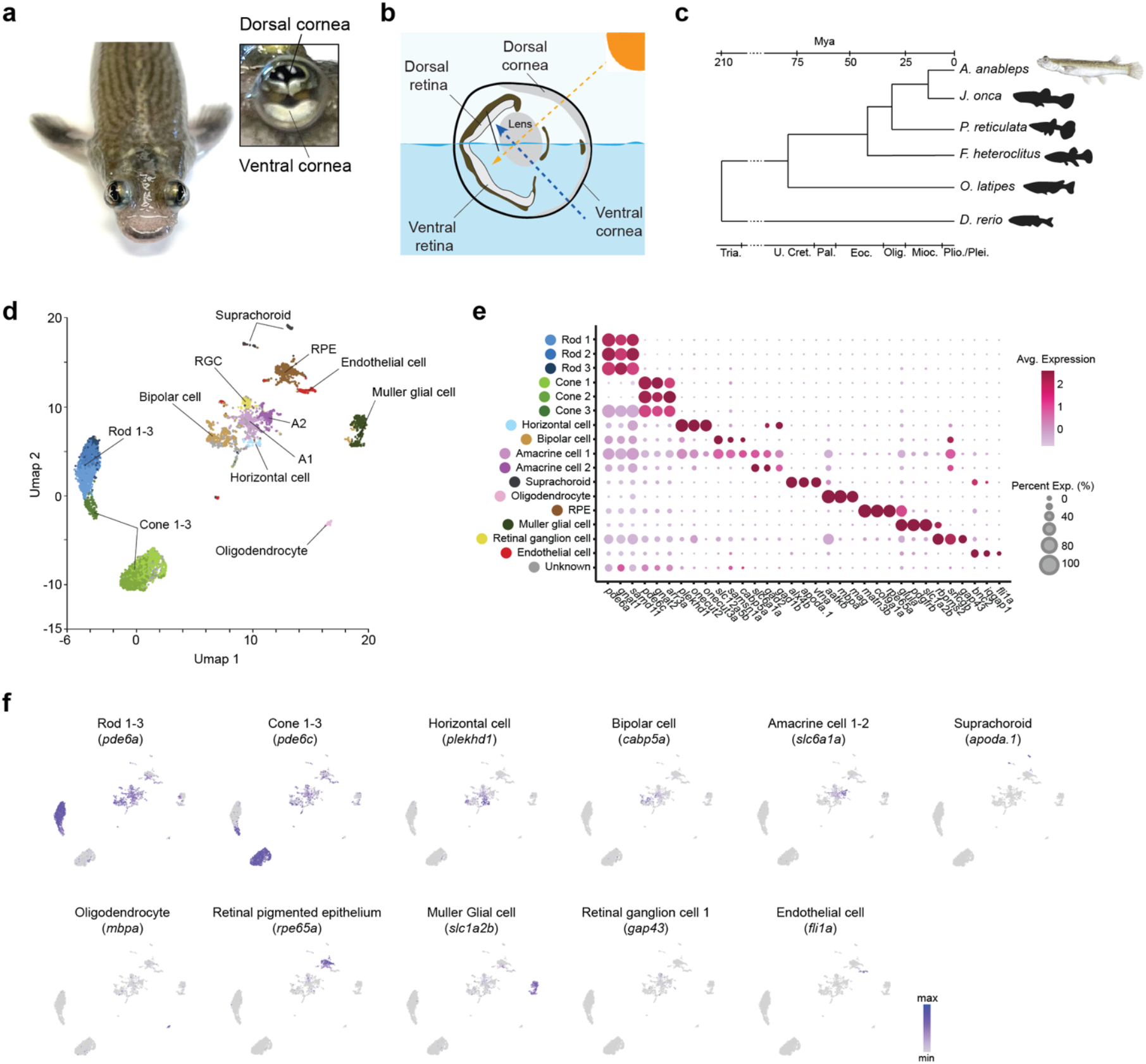
SnRNA-seq reveals cellular landscape of adult *Anableps* retina. **a**, Dorsal view of adult *Anableps* head, with an inset showing a lateral view of the eye, highlighting the dorsal and ventral corneas. **b**, Schematic representation of the visual field: aerial input (yellow arrow) through the dorsal cornea and aquatic input (blue arrow) through the ventral cornea. **c**, Phylogenetic relationship of *Anableps* relative to representative species of Anablepidae and other Cyprinodontiformes. **d**, UMAP plot shows automatic clustering of snRNA-seq gene expression profiles, revealing 17 distinct cell clusters; **e**, Dot plot showing expression of gene markers of retinal cell types identified in our dataset; **f**, UMAP plots show expression of representative gene markers for select retina cell types.

A previous survey of photoreceptor absorbance spectra in *Anableps* reported no difference in visual pigment composition in the dorsal and ventral retina^8^. However, subsequent *in situ* hybridization analyses suggested a more complex pattern, with ultraviolet (UV)- and short-wavelength-sensitive opsins distributed across the retina, middle-wavelength-sensitive cones confined to the ventral region, and long-wavelength-sensitive cones restricted to the dorsal region^9^. These seemingly conflicting findings have left the spatial organization and functional specialization of cone types in *Anableps* unresolved.

Anatomical studies have revealed that the *Anableps* ventral retina contains a denser ganglion cell population, consistent with enhanced sampling of the aerial visual field^10^. Similarly, histological analyses indicate that the ventral retina is thicker than the dorsal retina, suggesting a higher overall cell density in this region^11^. Furthermore, the middle wavelength opsin encoded by *opn1mw1* gene was found to be expressed in the entire ventral half of the retina^9^. Therefore, a central question is whether the putatively enlarged ventral half of the *Anableps* retina arose through an expansion of ventrally specified cell types, or through functional reprogramming of developmentally dorsal retinal domains. In the first scenario, ventral cell identities defined by canonical ventral patterning markers would have expanded to occupy a disproportionate portion of the retina. Alternatively, the dorsal retina, while retaining its molecular dorsal identity, may have acquired opsin expression profiles and photoreceptor specializations suited for aerial light detection. Distinguishing between these possibilities is essential to understanding how developmental patterning and photoreceptor diversification played a role in producing this evolutionary novelty.

Here, we use a multi-omics approach to investigate how the *Anableps* retina was remodeled to detect and process simultaneous aerial and aquatic light stimuli. Using single-nucleus RNA sequencing (snRNA-seq), we identified the major retinal cell types comprising the *Anableps* retina and their molecular markers. Spatial transcriptomic profiling (spatial RNA-seq) of the *Anableps* eye allowed us to map gene expression profiles to their respective anatomical locations. Comparative gene synteny analyses confirmed that the *Anableps* opsin gene repertoire was already present in the common ancestor it shares with guppies. Interestingly, whereas the ventral retina of zebrafish contains cone types tuned to downwelling aquatic light, the *Anableps* ventral retina contains cones spanning all wavelength classes (UV, blue, green, and red), potentially enabling broad-spectrum sensitivity to aerial illumination.

Furthermore, expression analysis of dorsal-ventral patterning markers shows that ventral-type cells predominate in the *Anableps* retina, suggesting an expansion of cells specialized for aerial light processing. Assessment of opsin spectral tuning using OPTICS^12^ revealed that the red-shifted *Anableps opn1mw2* likely enables long-wavelength detection by *thrb*^+^ red cones in the ventral retina. In addition, we show via spatial RNA-seq how the dorsal and ventral retinal domains are established during eye development and maintained in adulthood. Finally, comparison of *Anableps* and killifish spatial RNA-seq datasets indicates a dorsal shift in the retina dorsal-ventral boundary in *Anableps*, effectively enlarging the ventral retinal territory. Together, these findings demonstrate that *Anableps* evolved an expanded and spectrally diversified ventral retina as an adaptation for aerial vision, a derived state contrasting sharply with the red-biased, water-adapted ventral retina of typical teleosts.

## Results

### SnRNA sequencing reveals retinal cell diversity in adult *Anableps*

To identify the cellular composition and transcriptional profiles of the *Anableps* retina, we performed snRNA-seq using a combinatorial barcoding approach (Parse Biosciences, 2024). Retinal nuclei were isolated from two technical replicates of whole adult retinas. Trailmaker (Parse Biosciences, 2024) was used to assess the quality control metrics, preprocess the data, and complete our snRNA-seq analysis. After preprocessing, we retained 5,668 high-quality nuclei for downstream analyses (Supplementary Fig. 1a). Unbiased clustering from all cells from our snRNA-seq data identified seventeen transcriptionally distinct cell populations (clusters) visualized using uniform manifold approximation and projection (UMAP) (Fig. 1d, Supplementary Fig. 1b). Cluster annotation based on transcriptionally distinct profiles of canonical retinal gene markers identified sixteen clusters, including photoreceptors, interneurons, and non-neuronal supporting cells (Fig. 1d and e, Supplementary Fig. 1b). Among photoreceptor cells, we identified rod subtypes (*pde6a*^+^/*gnat1*^+^/*samd11*^+^) and cone subtypes (*pde6c*^+^/*gnat2*^+^/*arr3a*^+^) (Fig. 1d,e). The retinal interneuron cells included a horizontal cell cluster (*plekhd1*^+^/*onecut2*^+^/*onecut3a*^+^), a bipolar cell cluster (*slc12a5b*^+^/*samsn1a*^+^/*cabp5a*^+^), and two amacrine cell clusters (*slc6a1a*^+^/*gad2*^+^/*gad1b*^+^). The other clusters were characterized by suprachoroid cells (*alx4b*^+^/*apoda*.*1*^+^/*vtna*^+^), oligodendrocytes (*aatkb*^+^/*mbpa*^+^/*mag*^+^), retinal pigment epithelium (RPE) (*matn3b*^+^/*col9a1a*^+^/*rpe65a*^+^), Müller glia (*glula*^+^/*pdgfrb*^+^/*slc1a2b*^+^), retinal ganglion cells (RGC) (*rbpms2*^+^/*sncgb*^+^/*gap43*^+^), and endothelial cells (*bnc2*^+^/*iqgap1*^+^/*fli1a*^+^) (Fig. 1d,e, Supplementary Fig. 1b). One cluster remained unclassified due to an ambiguous gene expression profile (Fig. 1d,e, Supplementary Fig. 1b). UMAP plots showed gene marker expression within expected cell populations, including rods (*pde6a*), cones (*pde6c*), horizontal cells (*plekhd1*), bipolar cells (*cabp5a*), amacrine cells (*slc6a1a*), suprachoroid (*apoda*.*1*), oligodendrocytes (*mbpa*), RPE (*rpe65a*), Müller glia cells (*slc1a2b*), retinal ganglion cells (*gap43*), and endothelial cells (*fli1a*) (Fig. 1f). The identification of multiple photoreceptor subtypes, diverse interneuron populations, and major supporting cell classes provided a comprehensive cellular map of the adult retina of *Anableps* at single-cell resolution for downstream analyses.

### Spatial gene expression domains of eye cell types

Our snRNA-seq analysis revealed seventeen transcriptionally distinct retinal cell clusters. To determine how these molecularly defined cell types are spatially arranged within the *Anableps* retina, we conducted spatial RNA-seq of the whole adult *Anableps* eye (Fig. 2a,b) using the Visium platform (10x Genomics). Unsupervised k-means clustering (k = 6) using Loupe Browser v8.1.2 (10x Genomics) identified transcriptionally distinct clusters, and assessment of the top 10 genes differentially expressed in each cluster identified characteristic marker genes (Fig. 2c).

**Fig. 2.**
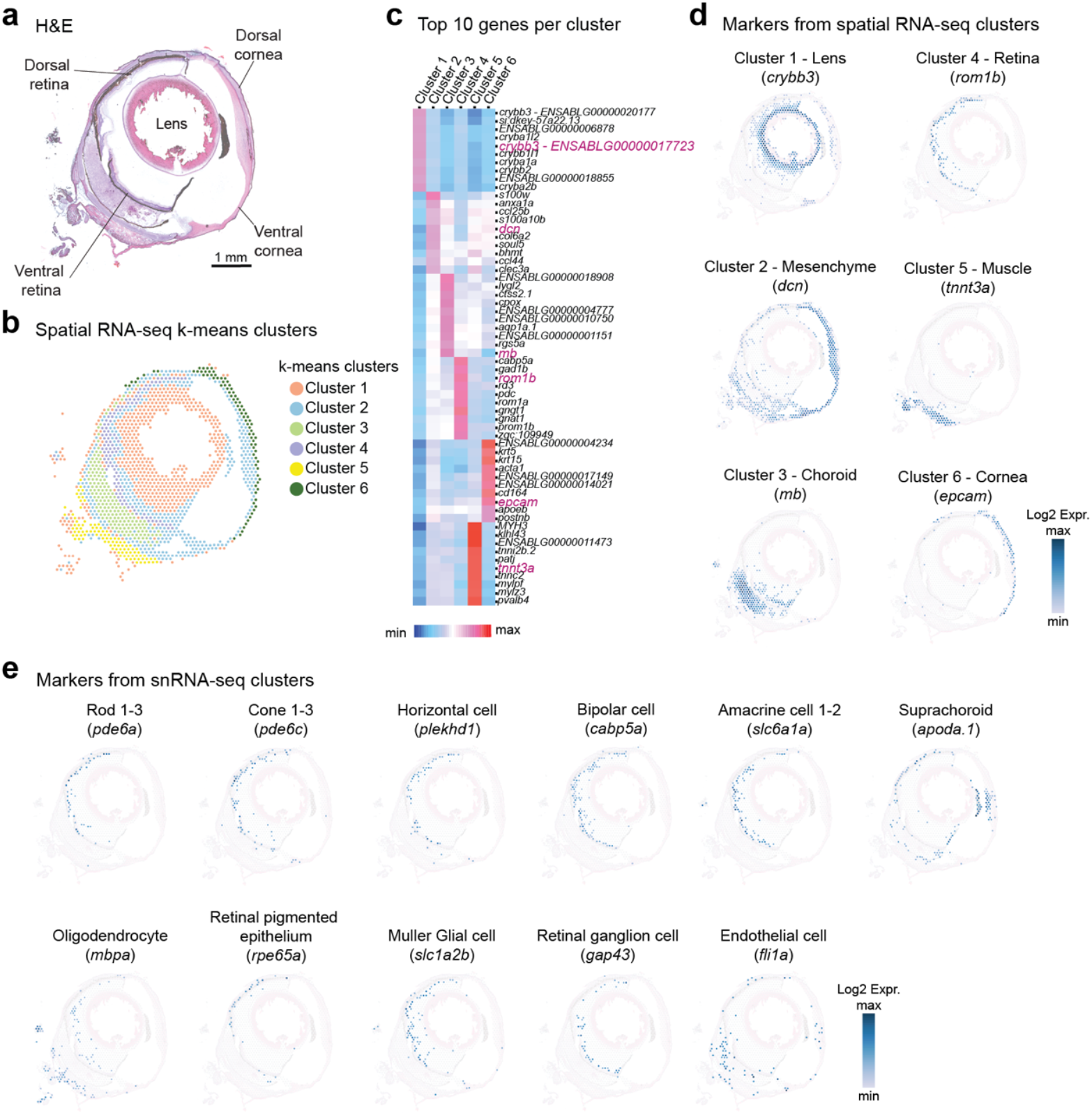
Spatial transcriptomics profiling of the adult *Anableps* eye. **a**, Histological section used for spatial RNA-seq of the *Anableps* eye. **b**, Unbiased k-means clusters identify six different gene expression territories in the *Anableps* eye. **c**, Heatmap showing top 10 differentially expressed genes per cluster. **d**, Eye genes marker from spatial RNA-seq clusters: *crybb3* (lens), *dcn* (mesenchymal), *mb* (choroid), *rom1b* (retina), *tnnt3a* (muscle), and *epcam* (cornea). **e**, Retina gene markers from our snRNA-seq clusters: *pde6a* (rod 1-3), *pde6c* (cone 1-3), *plekhd1* (horizontal cell), *canp5a* (bipolar cell), *slc6a1a* (amacrine cell 1-2), *apoda*.*1* (suprachoroid), *mbpa* (oligodendrocyte), *rpe65a* (retinal pigment epithelium), *slc1a2b* (muller glial cell), *gap43* (retinal ganglion cell), and *fli1a* (endothelial cell).

Based on these markers we assigned clusters to their likely tissue identities: lens (*crybb3*), mesenchyme (*dcn*), choroid (*mb*), retina (*rom1b*), muscle (*tnnt3a*), and cornea (*epcam*) (Fig. 2b–d). This spatial RNA-seq atlas established a framework for mapping additional cell-type-specific markers across the *Anableps* eye.

To validate that the retinal cell-type markers identified in our snRNA-seq analysis corresponded to their expected anatomical locations, we examined their spatial distribution within the *Anableps* eye. Although 10x Genomics Visium v1 slide architecture does not achieve single-cell resolution, as each capture spot encompassing transcripts from multiple neighboring cells (approximately 5-10 cells of the *Anableps*), the dataset nonetheless enabled robust localization of major retinal cell types across tissue layers. Expression of *pde6a* (rod 1–3), *pde6c* (cone 1–3), *plekhd1* (horizontal cell), *cabp5a* (bipolar cell), *slc6a1a* (amacrine cell 1-2), *rpe65a* (RPE), *slc1a2b* (Müller glia cell), and *gap43* (retinal ganglion cell) was confined to the retinal region, consistent with their known laminar organization. In contrast, *apoda*.*1* (suprachoroid) and *fli1a* (endothelial cell) were enriched in the choroid, whereas *mbpa* (oligodendrocyte) localized to the muscle underlying the sclera and adjacent mesenchyme (Fig. 2e). Together, these results confirm that the transcriptional identities inferred from our snRNA-seq data align with the expected spatial organization of retinal and extra-retinal tissues in the *Anableps* eye.

### Comparison of *Anableps* and zebrafish cone photoreceptor diversity and opsin gene repertoire

To compare cone photoreceptor diversity and opsin gene expression between *Anableps* and zebrafish, we subclustered cone cells from our *Anableps* snRNA-seq dataset and a publicly available single-cell RNA-seq (scRNA-seq) dataset from adult zebrafish retina^7^. Unsupervised subclustering identified seven transcriptionally distinct cone subtypes in both *Anableps* (Fig. 3a,b) and zebrafish (Fig. 3e,f; Supplementary Fig. 3). In *Anableps*, cone subtypes were defined by enriched expression of the following opsins: UV (*opn1sw1*^+^; SWS1), blue 1 and 2 (*opn1sw2_2*^+^; SWS2b), green 1 and 2 (*opn1mw1*^+^; RH2-2), red 1 and 2 cones. Red 1 expressed *opn1mw2* (RH2-1) while red 2 expressed two tandem copies of *opn1lw1* (LWS1), currently annotated in the *Anableps* genome (fAnaAna1) as a single transcript (here referred to as *opn1lw1_1&2)*, as well as *opn1lw2* (LWS2) (Fig. 3a,b). Similarly, zebrafish cones comprised UV (*opn1sw1*^+^; SWS1), blue 1 and 2 (*opn1sw2*^+^; SWS2), green 1 and 2 (*opn1mw1*^+^; RH2-1/*opn1mw2*^+^; RH2-2/*opn1mw3*^+^; RH2-3), and red 1 and 2 (*opn1lw1*^+^; LWS1/*opn1lw2*^+^; SWS2) cones (Fig. 3e,f). Cone subtype identities were validated by expression of established zebrafish marker genes^7,13^: *tbx2b* (UV), *hs3st3b1a* (blue), *fibcd1a* (green), and *thrb* (red) (Fig. 3b,f). Consistent with previous reports, *opn1mw4* (RH2-4) was expressed in a unique population of double cones previously classified as ventral retina green-red cones expressing high levels of *cycsb*^7^. Likewise, in agreement with previous findings^9^, we did not detect significant expression of *Anableps* opn1sw2_1 (SWS2a) or *opn1lw1_3* (Fig. 3b-d; Supplementary Fig. 3).

**Fig. 3.**
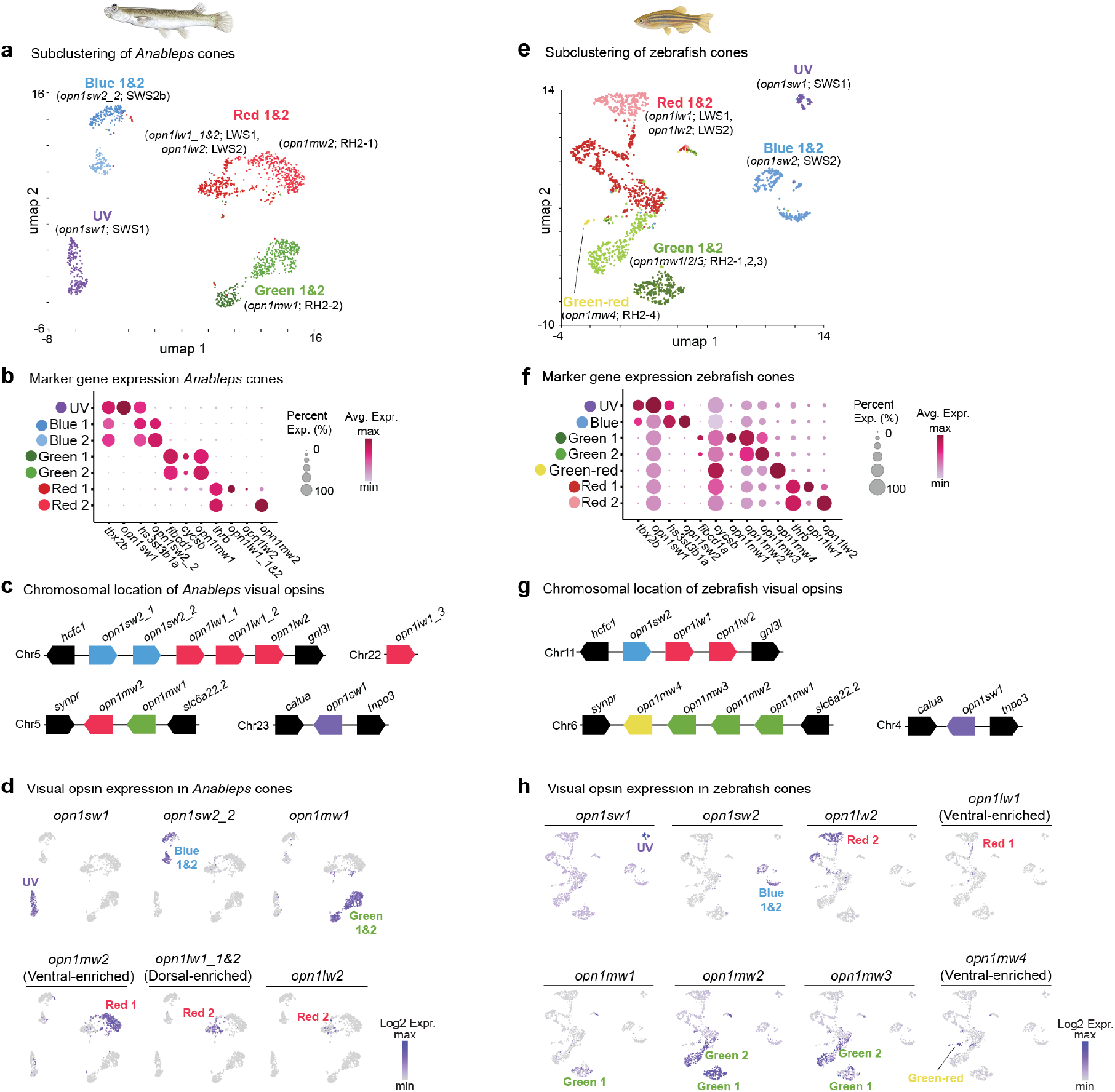
Comparison of cone cell diversity and opsin gene expression in *Anableps* and zebrafish retinas. **a, e**, UMAP plots showing the cone subclusters in *Anableps* (**a**), and in zebrafish (**e**). **b, f**, Dot plot of cone subtype markers and visual opsin genes in *Anableps* (**b**) and zebrafish (**f**). **c, g**, Chromosomal organization of *Anableps* (**c**) and zebrafish (**g**) visual opsin genes. **d, h**, UMAP plots showing the expression patterns of visual opsins in *Anableps* (**d**) and zebrafish (**h**) cones.

The availability of a high-quality *Anableps* genome (fAnaAna1) allowed us to re-examine the opsin gene repertoire and chromosomal organization (Fig. 3c,g). We confirmed nine visual opsin genes^9,14^ in *Anableps* genome, contrasting to the eight visual opsins of zebrafish^15^. Although the number of opsin paralogs in *Anableps* differs from zebrafish, their counterparts are present in the guppy and the one-sided livebearer^9,16^. Conserved synteny analyses confirmed that the *Anableps* opsin gene complement was already present in their common ancestor with the guppy and killifish (Supplementary Fig. 2a-c).

Finally, mapping visual opsin expression across cone subclusters revealed a clear correspondence between opsin identity and cone subtype in both species (Fig. 3d,h). In both zebrafish and *Anableps* each cone subtype formed two separated clusters, although this subdivision was more pronounced in *Anableps* (Fig. 3a,e). Notably, in *Anableps, opn1mw2* (ventral-enriched) and *opn1lw1_1&2* (dorsal-enriched), previously shown to exhibit asymmetric expression along the dorsal-ventral axis^9^, each distinctly marked one of the two red cone subtypes (Fig. 3d; Supplementary Fig. 3a). This organization supports the interpretation that the two red cone clusters in *Anableps* correspond to dorsally and ventrally localized photoreceptors

### Expanded cone diversity across the dorsal-ventral retina in *Anableps*

To assess how cone subtypes are distributed along the dorsal-ventral axis of the *Anableps* and zebrafish retinas, we classified cone cells according to established dorsal-ventral markers. Our *Anableps* snRNA-seq dataset captures nascent nuclear transcripts, which are generally less abundant than cytoplasmic mRNAs measured in scRNA-seq. This limits detection of lowly expressed transcription factors such as the dorsal marker *tbx5b* and ventral marker *vax2*^17^. We confirmed by hybridization chain reaction RNA fluorescence *in situ* hybridization (HCR RNA-FISH) that *Anableps tbx5b* and *vax2* are expressed in the dorsal and ventral regions of the retina, respectively, during development, and that these expression domains are maintained in the adult retina of both *Anableps* and zebrafish (Supplementary Fig 4).

To identify robustly expressed dorsal-ventral markers in *Anableps*, we searched for genes with the highest expression correlation with *tbx5b* and *vax2*. Among *tbx5b*-correlated genes, *cngb3*.*2*, which encodes the cyclic nucleotide-gated channel β3 subunit that account for ∼50% of autosomal recessive achromatopsia cases in humans^18^, showed the strongest expression, over 10-fold greater than that of *tbx5b* (Supplementary Fig. 5a). Conversely, *ephb3*, a well-established ventral retina marker in vertebrates^19,20^, showed an expression level approximately 3-fold higher than *vax2* and was the most highly expressed gene correlated with *vax2* (Supplementary Fig. 5b). UMAP and violin plots revealed *cngb3*.*2* enrichment in subsets of UV, blue 2, green 1 and red 1 cone clusters (Fig. 4a), whereas *ephb3* was enriched in complementary populations: *cngb3*.2-negative UV, blue 1, green 2 and red 2 cones (Fig. 4b). The two markers showed largely nonoverlapping expression across cone clusters, with nearly 60% of *Anableps* cone cells expressing the ventral marker *ephb3* (Fig. 5c).

**Fig. 4.**
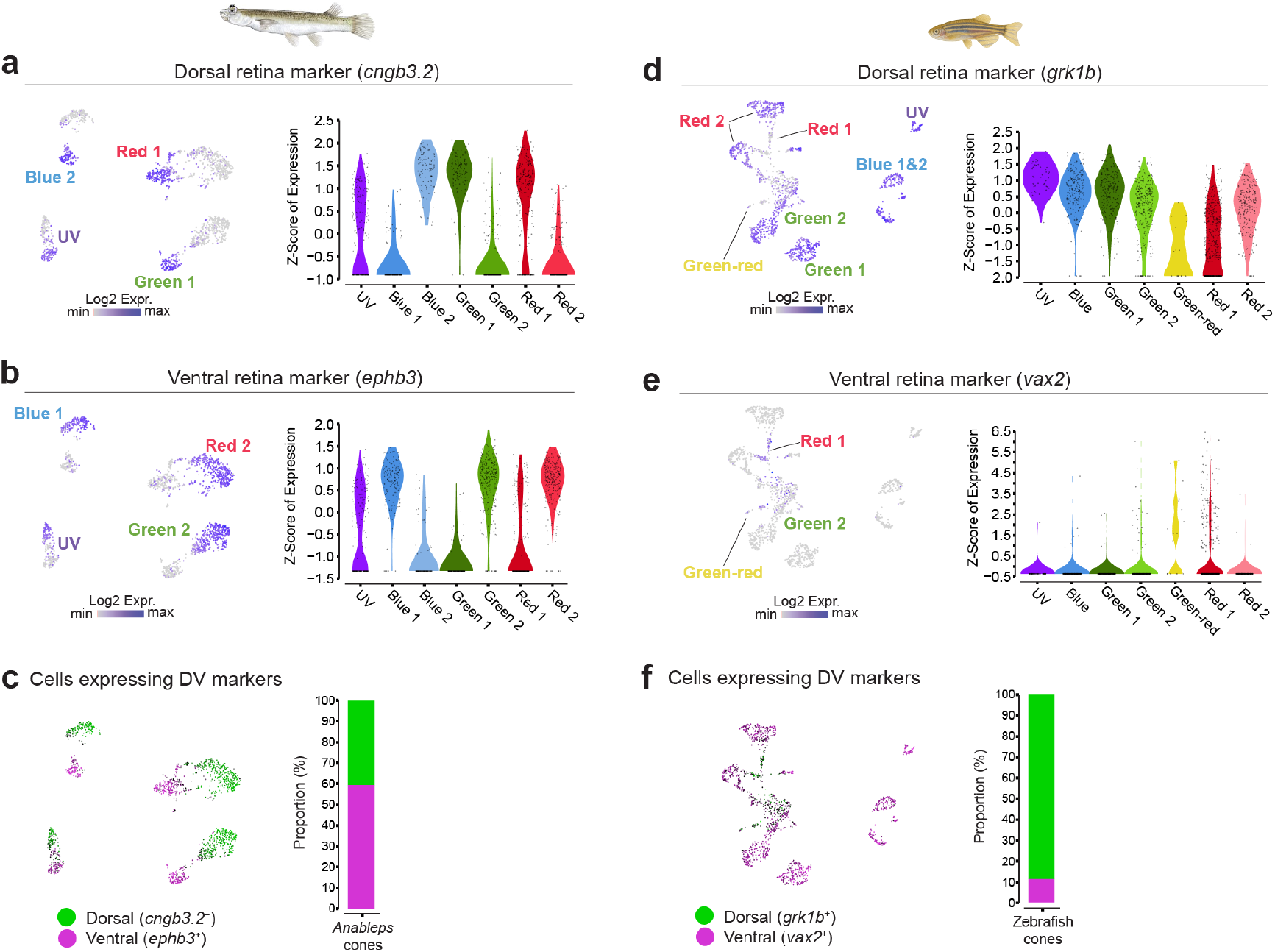
Cone diversity along the dorsal-ventral axis of the *Anableps* and zebrafish retinas. **a, d**, UMAP and dot plots showing expression of *cnhb3*.*2* and *grk1b* in the cone clusters of the *Anableps* (**a**) and zebrafish (**d**) retinas. **b, e**, UMAP and dot plots showing expression of *ephb3* and *vax2* in the cone clusters of the *Anableps* (**b**) and zebrafish (**e**) retinas. **c, f**, UMAP plots highlighting dorsal (green) and ventral (magenta) cells identified by dorsal-ventral marker expression in the *Anableps* (**c**) and zebrafish (**f**) retinas; bar plot showing the proportion of number of cells assigned to each retinal region based on DV marker expression.

**Fig. 5.**
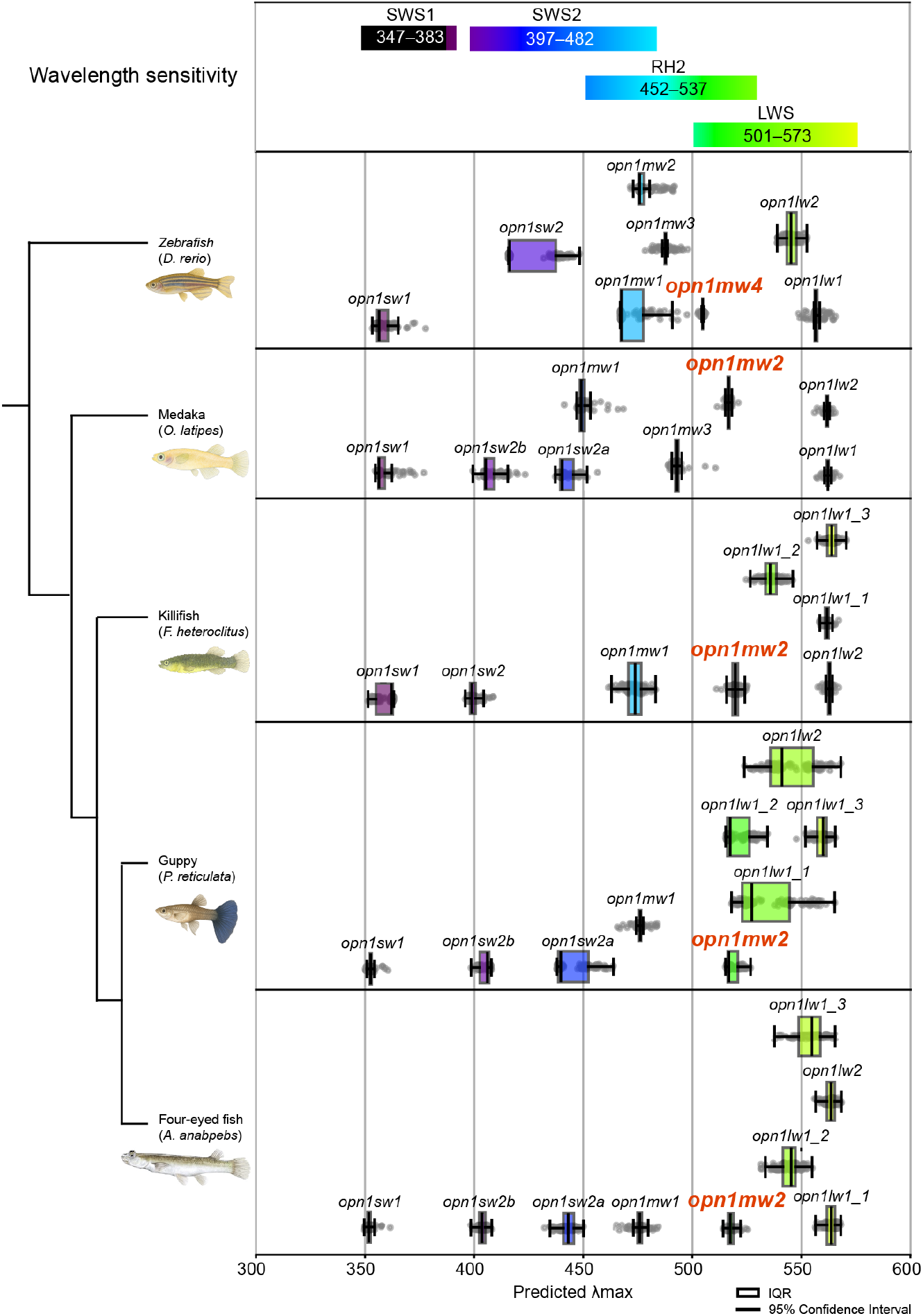
Prediction of opsin sensitivity spectra indicates a red shift of *opn1mw2* orthologs. Prediction of peak wavelength sensitivity (λmax) of visual opsins of zebrafish, medaka, killifish, guppy and *Anableps*. IQR, interquartile range.

In the zebrafish scRNA-seq dataset, we used *grk1b* (a recently identified dorsal cone marker^7^) and *vax2* as dorsal and ventral markers, respectively. UMAP and violin plots showed broad *grk1b* expression across UV, blue (1 and 2), green (1 and 2), and red 2 clusters, and partially in green-red and red 1 cone clusters (Fig. 4d). In contrast, *vax2* was enriched in red 1 cones and present in subsets of green-red and green 2 clusters (Fig. 4e). UMAP plot confirmed largely exclusive expression of *grk1b* and *vax2*, with *vax2*^+^ cells representing approximately 12% of all cones in the zebrafish retina (Fig. 4f).

Together, these findings indicate that in *Anableps*, both dorsal and ventral retina regions contain a full complement of cone subtypes, encompassing all wavelength classes. In contrast, zebrafish, consistent with previous reports^7,21^), show a dorsal bias toward short-wavelength (UV/blue/green) cones. These results suggest that during evolution, the *Anableps* ventral retina underwent a relative expansion in cone cell number and diversity compared to the dorsal retina.

### Spectral tuning and wavelength sensitivity of *Anableps* opsins

Our snRNA-seq dataset enabled the classification of cone subtypes according to markers associated with distinct wavelength sensitivities. In addition, analyses of dorsal-ventral patterning markers allowed us to assign each cone cluster to either dorsal or ventral retinal identity in both *Anableps* and zebrafish. Notably, *Anableps opn1mw2* and zebrafish *opn1mw4*, though members of the middle-wavelength-sensitive (MWS) opsin class, were both expressed in the ventral retina in *thrb*^+^ red cones in *Anableps* and in ventrally enriched green-red cones in the zebrafish (Fig. 3b,f). This suggests that these MWS opsins may have undergone red-shifted spectral tuning, allowing ventral photoreceptors to detect longer-wavelength light stimuli.

To estimate the peak wavelength sensitivity (λmax) of *Anableps* opsins, we applied OPTICS, a recently developed machine-learning framework that predicts photopigment absorbance spectra directly from opsin amino acid sequences^12^. The OPTICS approach first extracts features related to spectral tuning from each opsin sequence, then uses a neural network trained on a broad dataset of vertebrate opsins with known λmax values. The algorithm infers how variant amino acids drive shifts in wavelength sensitivity, enabling prediction of λmax even in species where no experimental pigment reconstitution has been performed.

To validate OPTICS predictions, we first analyzed zebrafish opsins whose λmax values have been experimentally characterized for short-wavelength (*opn1sw1*: ∼365 nm; *opn1sw2*: ∼416 nm)^22^, middle-wavelength (*opn1mw1–4*: 467, 476, 488, 505 nm, respectively)^15^ and long-wavelength (*opn1lw1*: 558 nm; *opn1lw2*: 548 nm)^15^ pigments. OPTICS predictions closely matched these values (*opn1sw1*: 355 nm; *opn1sw2*: 416 nm; *opn1mw1–4*: 467, 475, 487, 505 nm; *opn1lw1*: 557 nm; *opn1lw2*: 545 nm) (Fig. 5; Supplementary Data 1), demonstrating strong predictive accuracy. Consistent with prior studies, *opn1mw4*, which in zebrafish is the most red-shifted of the MWS opsins, was expressed in green-red cones enriched in the ventral retina (Fig. 3e,f,h)^7^.

Next, to explore the evolution of opsin spectral tuning in *Anableps*, we estimated λmax values for visual opsins from medaka (*O. latipes*), mummichog (*F. heteroclitus*), guppy (*P. reticulata*) and *Anableps*. Across these species, the most striking pattern was a red shift in *opn1mw2* orthologs, placing them within the range typical of long-wavelength (LWS) fish opsins^23^. Together, these findings indicate that the red shift of *opn1mw2* predates the emergence of aerial vision in *Anableps* and suggest that the *Anableps opn1mw2*, with an estimated λmax = 517 nm (Fig. 5; Supplementary Data 1), enables long-wavelength detection by *thrb*^+^ red cones in the ventral retina.

### Dorsal expansion of the ventral retina in *Anableps*

Previous reports^10,11^ and our current findings suggest that the ventral retina in *Anableps* has expanded in cell content relative to the dorsal retina. To examine the establishment of dorsal-ventral retinal territories during development and whether increased cell numbers correspond to spatial expansion, we performed spatial RNA-seq on developing *Anableps* eyes. Analysis of ventral markers (*ephb2b, ephb3, smoc1*, and *opn1mw2*) and dorsal markers (*prph, tbx5b, nr2f2* and *fgf24*) across developmental stages 3, 4 and 5 and in the adults revealed gradual regionalization of marker expression, which persisted into adulthood (Fig. 6a,b). Combined expression of ventral markers indicated that the ventral retina ultimately occupies roughly half of the dorsal-ventral length of the *Anableps* retina (Fig. 6a, bottom right). Next, we assessed the same dorsal-ventral marker set in the adult killifish eye using spatial RNA-seq (Fig. 6c,d). In the killifish, the ventral retina occupied about one-fourth to one-third of the dorsal-ventral retinal length (Fig. 6c). These findings support the conclusion that the ventral retina expanded dorsally in *Anableps* sometime after the divergence of Anablepidae and Fundulidae.

**Fig. 6.**
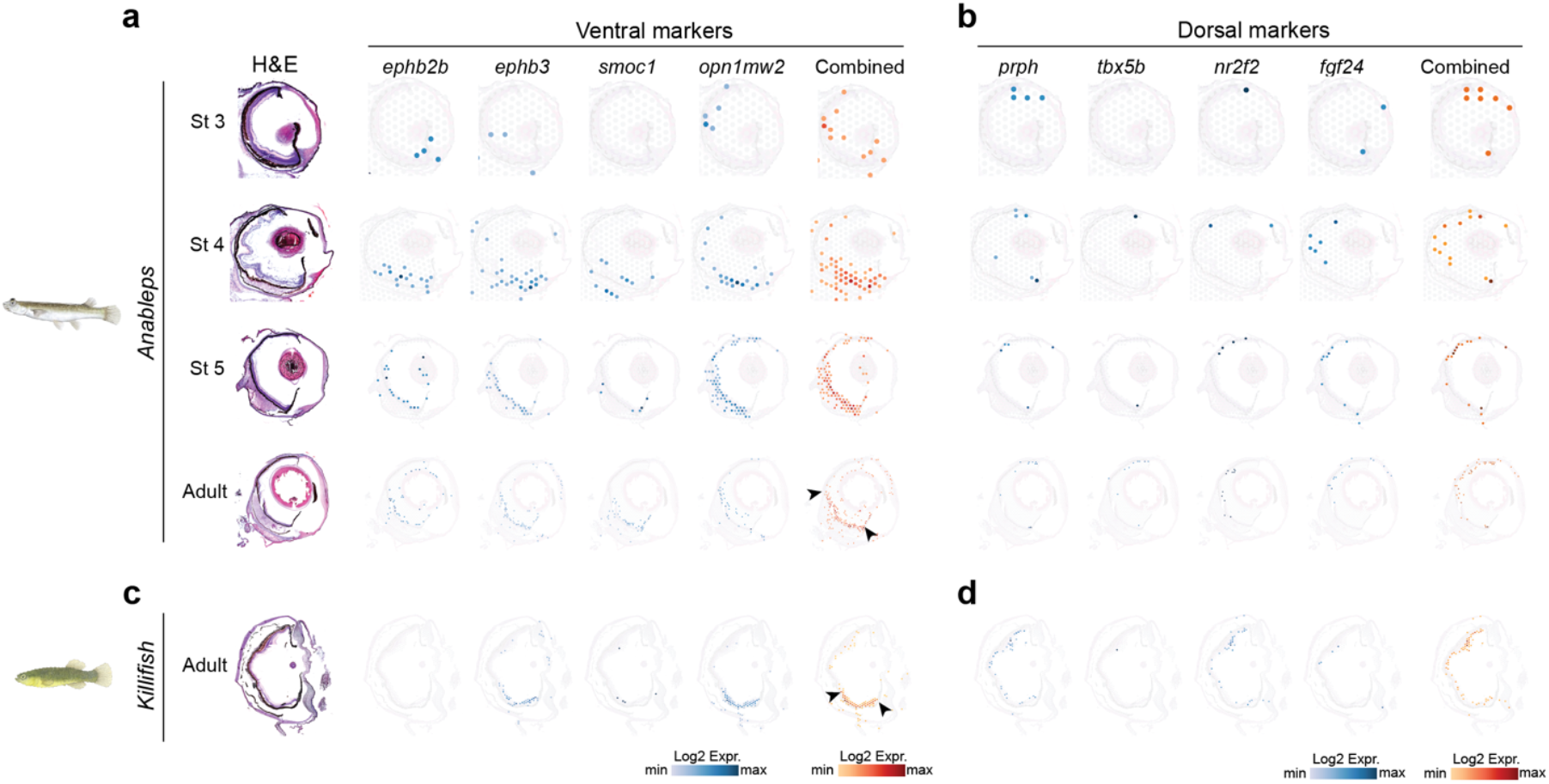
Comparative spatial transcriptomics of *Anableps* and killifish reveals dorsal expansion of the ventral retina. **a-c**, Spatial RNA-seq showing expression of ventral (***a*, c**) and dorsal (**b, d**) markers across developmental stages and adulthood in *Anableps* (**a, b**) and adult killifish (**c, d**). Combined expression of these markers highlights the expansion of the ventral retina in *Anableps* relative to killifish, indicated by black arrows in **a, c**.

**Fig. 7.**
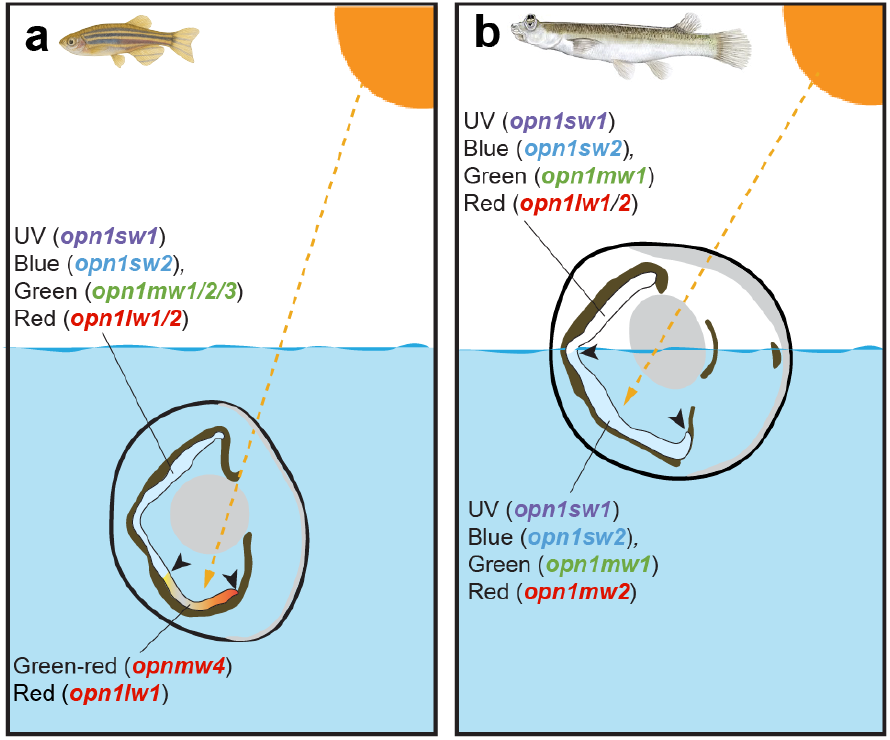
Retinal and opsin adaptations underlying dual-environment vision in *Anableps*. **a, b**, Schematic representation of opsin gene expression and cone type composition of dorsal and ventral retina regions in zebrafish (**a**) and *Anableps* (**b**) eye. Black arrowheads indicate approximate region occupied by the ventral retina.

## Discussion

Morphological innovations allow organisms to adapt to new niches and exploit new ecological opportunities, yet how such innovations arise has been a longstanding problem in evolutionary biology. A mechanistic understanding of the cellular and molecular bases of morphological innovations relies on organismal models amenable to experiments in a laboratory setting. Here we leveraged cutting-edge transcriptomic techniques to survey the *Anableps* retina at single cell resolution and obtain a comprehensive, spatially resolved map of the developing and adult retinas, and employed machine-learning based approach to evaluate evolution of *Anableps* opsin gene sensitivity. Furthermore, our comparative analysis to an existing zebrafish scRNA-seq^7^ and our newly generated killifish spatial RNA-seq datasets provided evolutionary context to the unique adaptations that led to the remodeling of the *Anableps* ventral retina.

In fully aquatic teleosts, the ventral-most retina is classically adapted to monitor downwelling light, which comes from above water and penetrates the aquatic column^24^. Because water preferentially attenuates longer wavelengths (reds), downward light tends to be blue-shifted or green-biased. Thus, the ventral retina often expresses opsins and cone types tuned to shorter to middle wavelengths (e.g. SWS1, SWS2, RH2), sometimes with reduced or absent long-wavelength (LWS) cones. However, *Anableps* occupies a strikingly different niche: the air-water interface. As a result, its ventral retina is exposed to aerial (above-water) light. Aerial light is richer in long wavelengths (reds, oranges) and broader spectrum overall, compared to downwelling underwater light^25^. Hence, the ventral retina in *Anableps* is expected to be under selection to respond more broadly to the full visible spectrum. Concordantly, we showed that the *Anableps* contains the full complement of cone types spanning all wavelength classes. Although the *Anableps* ventral cone do not express traditional LWS genes, our results provide evidence that the red-shifted *opn1mw2* may fulfill the task of LWS opsins in *Anableps* ventral retina cells. Interestingly, this shift of *opn1mw2* to LWS sensitivity spectrum is also predicted in the medaka, killifish, and guppy, indicating that this feature evolved before *Anableps* adaptation to the water-air interface.

Because the ventral retina captures a larger fraction of the visual field above water, selection would be expected to favor both its expansion and diversification of cone types. Indeed, several anatomical and molecular observations support this notion: the ventral half of the *Anableps* retina contains approximately 3.6-fold more neurons in the ganglion cell layer than the dorsal half^10^, the ventral inner nuclear layer is roughly twice as thick^11^, and *opn1mw2* expression is confined to the ventral retina with a sharp boundary at the dorsal-ventral midline^9^. Collectively, these findings point to a substantial enlargement and functional specialization of the ventral retinal territory in *Anableps*.

An unresolved question is the developmental origin of this expansion. The disproportionate ventralization of the retina could have arisen either through a true enlargement of ventrally specified territories during development or through functional reprogramming of regions that were originally dorsal in identity. To distinguish between these possibilities, we examined the expression of canonical dorsal-ventral markers during *Anableps* development and in the adult retina, alongside comparisons to the adult killifish. Our analyses show that the ventral half of the *Anableps* retina retains a ventral molecular identity, as indicated by persistent expression of ventral markers. Thus, the extended *opn1mw2* domain reflects an expansion of ventrally specified tissue rather than dorsal cells acquiring ventral-like opsin expression. This interpretation is further supported by the expression killifish *opn1mw2* ortholog, which is likewise confined to the ventralmost region of the retina within the *ephb3*-positive domain.

Together, our findings illuminate how developmental patterning, opsin diversification, and domain-specific expansion were integrated to generate the distinctive visual system of *Anableps*. More broadly, these results exemplify how the modification of ancestral developmental programs can give rise to new sensory architectures and ecological capabilities, highlighting a general route by which morphological novelties emerge during vertebrate evolution.

## Material and Methods

### Animal work

*Anableps* and zebrafish were maintained and used in accordance with approved Louisiana State University (LSU) IACUC protocols IACUCAM-25-094 and IACUCAM-24-003, respectively. Fish were obtained from commercial vendors. *Anableps* were maintained in colonies of 8–10 individuals in large tanks within a recirculating freshwater system at 27-28 °C and 12 h light/12 h dark cycle. Zebrafish were kept in small tanks within a recirculating freshwater system at 27-28 °C and a 12 h light/12 h dark cycle. Prior to eye dissection, animals were euthanized in 0.3 % MS-222.

### Nuclei preparation and snRNA-seq sequencing

A pair of retinas from adult *Anableps* specimen was dissected and minced with a razor blade in a petri dish on ice. The minced tissue was resuspended in 1 mL of lysis buffer (ATAC-seq lysis buffer, Active Motif Inc.) and then homogenized in a pre-chilled Dounce homogenizer (3 strokes). The homogenate was filtered through a 40 µm cell strainer and immediately centrifuged in an Eppendorf Protein LoBind tube at 500 xg at 4 °C for 5 minutes in a swinging bucket rotor. The pellet was quickly washed with 1 mL 1X PBS/0.5 % BSA and then resuspended in 1 mL of 1X PBS/0.5 % BSA. Nuclei were counted using a hemocytometer and then a volume corresponding to 10^6^ nuclei was transferred to a new tube and centrifuged at 500 xg at 4 °C for 5 minutes. Nuclei were fixed and frozen using the Evercode Nuclei Fixation Kit v3 (Parse Biosciences) following the manufacturer’s protocol. Two technical replicates were prepared. Fixed and frozen nuclei were then processed to generate two RNA-sequence libraries aiming for 5,000 nuclei in each of them, using the Evercode WT Mini v3 Kit following the manufacturer’s protocol.

### SnRNA-seq processing and downstream analysis

SnRNA-seq FASTQ files were processed using the Trailmaker^TM^ piping module pipeline (https://app.trailmaker.parsebiosciences.com/; v1.4.0, Parse Biosciences, 2024). Heatmaps, dot plot, violin plots, and UMAP dimensionality reduction plots were generated using the Seurat v5 R package in R Studio and the TrailmakerTM platform. ScRNA-seq data from zebrafish was obtained from the Gene Expression Omnibus (GEO; accession number GSE175929)^7^. The data was reprocessed using Cell Ranger (10x Genomics). Cellranger mkref was used to index the zebrafish genome (GRCz11), and cellranger count was employed to map the reads to the reference genome. After processing, the output files (barcodes.tsv.gz, features.tsv.gz, and matrix.mtx.gz) were uploaded to Trailmaker^TM^, where the subsequent analysis was performed. A small subset of cone-annotated cells in the *Anableps* dataset expressed rho transcripts. Because co-expression of rod and cone markers con result doublets or ambient RNA contamination rather than a distinct biological population, these cells were excluded from further interpretation.

### Spatial RNA-seq libraries preparation and sequencing

Spatial transcriptomics was performed using the Visium Spatial Gene Expression platform (10X Genomics). The eyes of *Anableps* and killifish were processed following the Visual Spatial Protocol Tissue Preparation from 10X Genomics. Briefly, *Anableps* (adult, stage 3, stage 4, and stage 5) and adult killifish were euthanized using 0.3 % MS-222, the eyes (adults and stage 5) were removed, and the heads (stage 3 and 4) were flash-frozen in Tissue-Tek Optimal Cutting Temperature (OCT) compound (Tissue Tek Sakura) using liquid nitrogen and isopentane, then stored at −80 °C. The tissues were cryosectioned at 10 µm at −20 °C using a Leica CM1520 cryostat and placed onto Visium Spatial Gene Expression slides (10X Genomics). The slides were fixed in cold methanol and stained with Hematoxylin and Eosin (H&E) (Sigma-Aldrich). Tissue permeabilization and reverse transcription (RT) were performed using the Visium Spatial Tissue Optimization Kit (10X Genomics), with a permeabilization time of 18 minutes. Images of H&E-stained sections and tissue optimization were captured using a Leica DM6B Fluorescence Microscope, equipped with a Leica DFC450 color CCD and Hamamatsu sCMOS camera, using a 20x objective. Gene expression slides and reagent kits were prepared according to the 10X Genomics protocol to produce sequencing libraries. The libraries were sequenced by Novogene Corporation Inc. on a NextSeq 6000 (Illumina) platform, generating 80 million paired-end reads of 150 bp and 50,000 transcripts per spot.

### Spatial transcriptomics data processing and downstream analysis

Gene expression quantification for the Visium dataset was performed using Space Ranger (version 2.1.1, 10x Genomics). The *Anableps* genome (GCA_014839685.1) and killifish genome (GCF_011125445.2) were indexed with mkref under Space Ranger. Subsequently, reads were aligned and quantified using Space Ranger count. Clustering and gene analysis were conducted using the Loupe Browser from the 10X Genomics Visium platform (version 8.1.2).

### Machine learning prediction of opsin spectral sensitivity

Maximum wavelength sensitivity of visual opsins was predicted using the Opsin Phenotype Tool for Inference of Color Sensitivity (OPTICS), a machine learning approach that infers maximum wavelength sensitive (λmax) directly from amino acid sequences^12^. Each opsin sequence was aligned to the wildtype-vert-mnm reference database using MAFFT within the optics_predictions.py pipeline, with bootstrapping and BLASTp analyses performed using bovine opsin as a reference. Predicted λmax values were compared across species to assess spectral tuning shifts and interpret cone subtype expression patterns in *<u>Anableps</u>*. Protein accession numbers for each opsin were as follows: *D. rerio*: *opn1sw1* (NP_571394.1), *opn1sw2* (NP_571267.1), *opn1mw4* (NP_571329.1), *opn1mw3* (NP_878312.1),*opn1mw2* (NP_878311.1), *opn1mw1* (NP_571328.2), *opn1lw1* (NP_001300644.1), and *opn1lw2* (NP_001002443.1). *O. latipes*: *opn1sw1* (NM_001104656.1), *opn1sw2_1* (XM_020703465.2), *opn1sw2_2* (NM_001104654.1), *opn1lw1* (XM_004069094.4), *opn1lw2* (NM_001104694.1), *opn1mw3* (XM_004086132.4), *opn1mw2* (XM_004086130.4), and *opn1mw1* (NM_001104655.1). *F. heteroclitus*: *opn1sw1* (XM_036149145.1), *opn1sw2* (XM_036151441.1), *opn1lw1_1* (XM_036151440.1), *opn1lw1_2* (XM_012854938.3), *opn1lw1_3* (XM_012869094.3), *opn1lw2* (XM_036151579.1), *opn1mw1* (XM_012872728.3), and *opn1mw2* (XM_012859940.3). *P. reticulata*: *opn1sw1* (NM_001297490.1), *opn1sw2_1* (NM_001297456.1), *opn1sw2_2* (NM_001297454.1), *opn1mw1* (NM_001297458.1), *opn1mw2* (NM_001297460.1), *opn1lw1_1* (NM_001297452.1), *opn1lw1_2* (XM_008410572.2), *opn1lw1_3* (XM_008397471.2), and *opn1lw2* (NM_001297450.1). *A. anableps*: *opn1sw1* (ENSABLT00000040448.1), *opn1sw2_1* (ENSABLT00000001542.1), *opn1sw2_2* (ENSABLT00000001545.1), *opn1mw2* (ENSABLT00000017435.1), *opn1mw1* (ENSABLT00000017442.1), *opn1lw1_1* (ENSABLT00000001559.1), *opn1lw1_2* (ENSABLT00000001564.1), *opn1lw2* (ENSABLT00000001632.1), and *opn1lw1_1* (ENSABLT00000032578.1).

### Cryosectioning

Dissected eyes from adult and developmental stages of *Anableps* and zebrafish were flash-frozen using isopentane cooled with liquid nitrogen. Isopentane was dispensed into a metal container and placed in liquid nitrogen to equilibrate for about 10 minutes. Tissues were embedded in Tissue-Tek O.C.T. compound (Sakura) using appropriate molds for 30 minutes. Using forceps, the molds were placed into the equilibrated isopentane until frozen. Samples were stored at −80 °C for sectioning. The tissues were cryosectioned at 10 µm (thickness) at −20 °C using a Leica CM1520 cryostat. And the slides were stored at −80 °C until further use.

### HCR™ Gold RNA-FISH

Probes targeting *tbx5b* and *vax2* for HCR RNA-FISH were designed and synthesized as oligo pools by Molecular Instruments, for zebrafish and *Anableps*. HCR was performed on tissue sections using the HCR™ Gold RNA-FISH kit (Molecular Instruments) according to the manufacturer’s instructions. Briefly, sections were fixed in ice-cold 4% paraformaldehyde for 15 min at 4 °C, dehydrated in 50%, 70%, and two changes of 100% ethanol (5 min each, room temperature), rinsed in 1× PBS, and surrounded with a hydrophobic barrier. Slides were pre-hybridized for 10 min at 37 °C in HCR™ HiFi Probe Hybridization Buffer, then incubated overnight at 37 °C with 50–100 µl probe solution (probes diluted to 100 µl per slide in HCR™ HiFi Probe Hybridization Buffer). Post-hybridization, slides were washed four times for 15 min each in pre-warmed 1× HCR™ HiFi Probe Wash Buffer, pre-amplified for 30 min in HCR™ Gold Amplifier Buffer, and incubated overnight at room temperature with 100 µl of amplification solution containing 2 µl each of H1 and H2 hairpins denatured at 95 °C for 90 s and cooled for 30 min. Finally, sections were washed four times for 15 min each in 1× HCR™ Gold Amplifier Wash Buffer, mounted in antifade medium (with optional DAPI), coverslipped, and stored at 4 °C protected from light until imaging.

## Data availability

Raw sequencing data for single-nuclei RNA-seq and spatial transcriptomics RNA-seq generated in this study are deposited in the Gene Expression Omnibus (GEO) under the accession numbers: GSE306228 (single-nucleus RNA-seq data) and GSE306418 (Spatial Transcriptomics). Processed snRNA-seq data are available at the Broad Single Cell Portal (https://singlecell.broadinstitute.org/single_cell) (SCP: SCP3248). All code used in this study is available at our GitHub repository: https://github.com/lnperez90/Perez-et-al-2025

## Acknowledgments

We thank Igor Schneider for helpful feedback and comments. This work was funded by start-up funds from Louisiana State University (P.S.), and an NSF-Integrative Organismal Systems (IOS) grant (2409931, P.S.).

## Author Contributions

L.P., J.F.S., and P.S. designed research; L.P., L.S., A.D., K.P., C.S., J.F.S., and P.S. performed research; L.P., J.F.S., and P.S. analyzed data; J.F.S., and P.S. wrote the paper with input from all authors.

## Competing Interest Statement

The authors declare no competing interest.

## Inclusion & ethics statement

This research aligns with the inclusion & ethical guidelines embraced by Nature Communications.

## Supplementary Information

**Supplementary Fig. 1.**
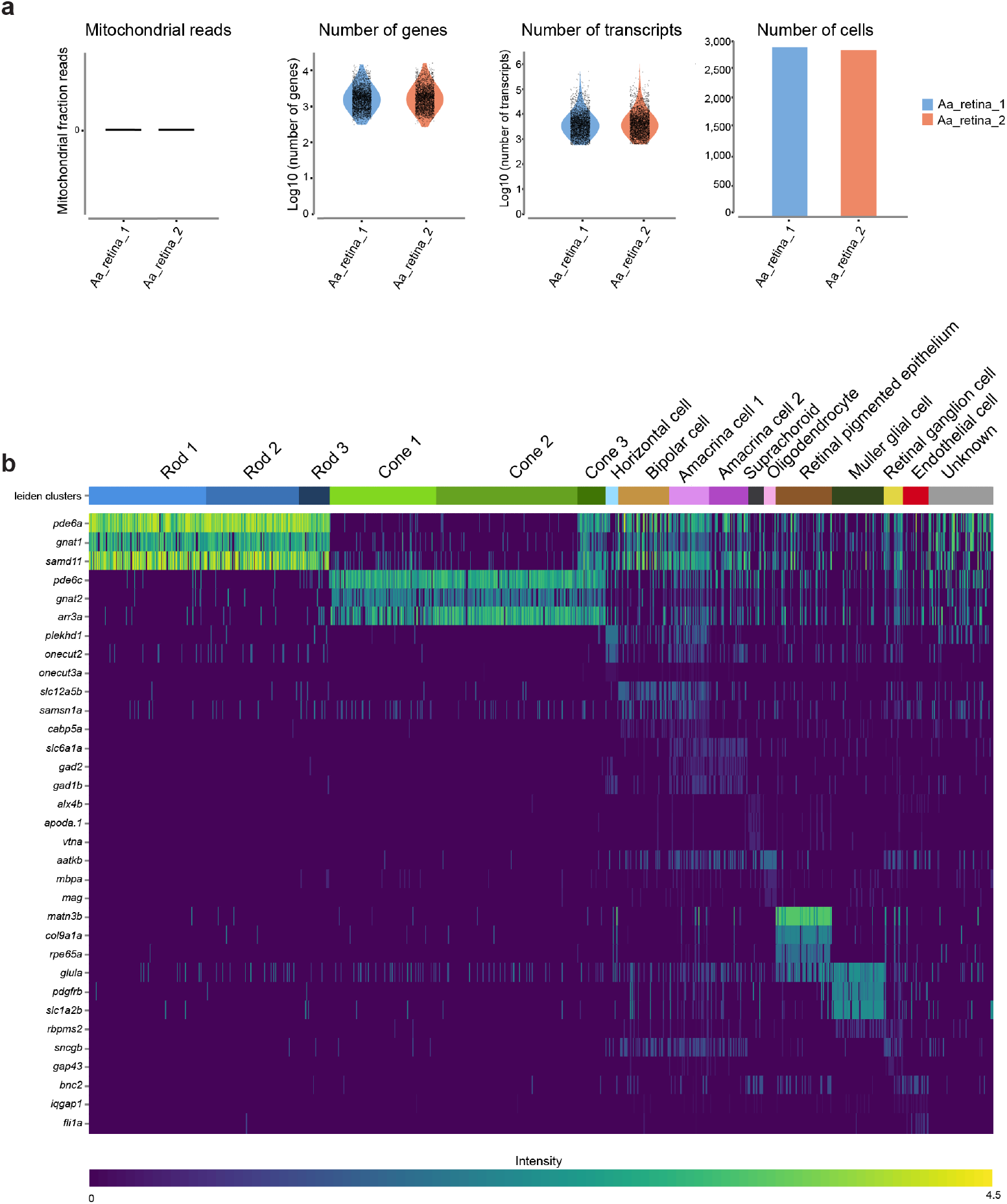
*Anableps* snRNA-seq quality control and marker gene expression across retina cell clusters. **a**, QC metrics showing mitochondrial read fraction, number of genes, transcripts, and cells (nuclei) across technical replicates. **b**, Heatmap showing gene markers expression across retina cell clusters.

**Supplementary Fig. 2.**
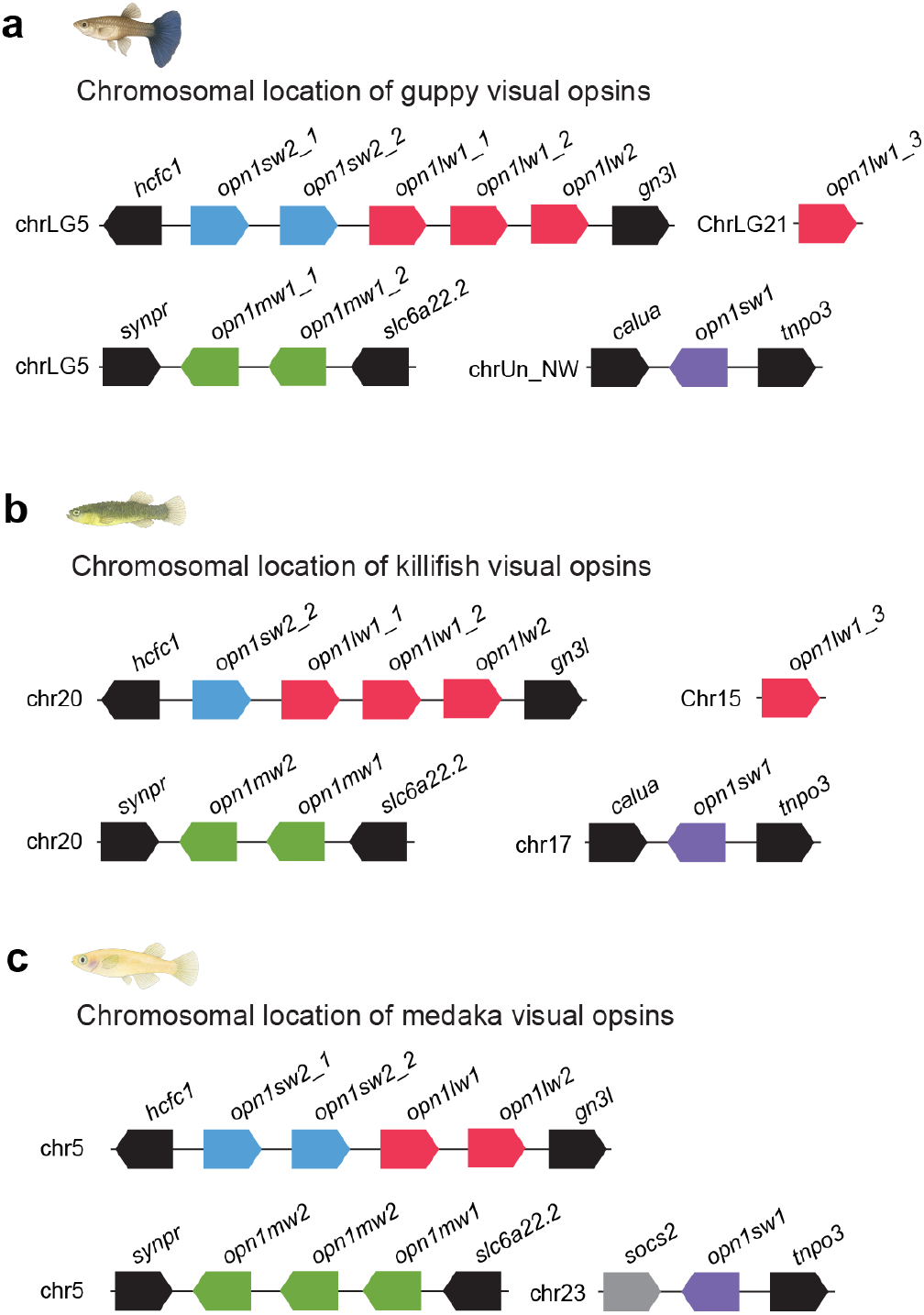
Gene organization of visual opsins in the genomes of guppy, killifish and medaka. Schematic drawings of genomic loci harboring visual opsin genes in guppy (**a**), killifish (**b**), and medaka (**c**).

**Supplementary Fig. 3.**
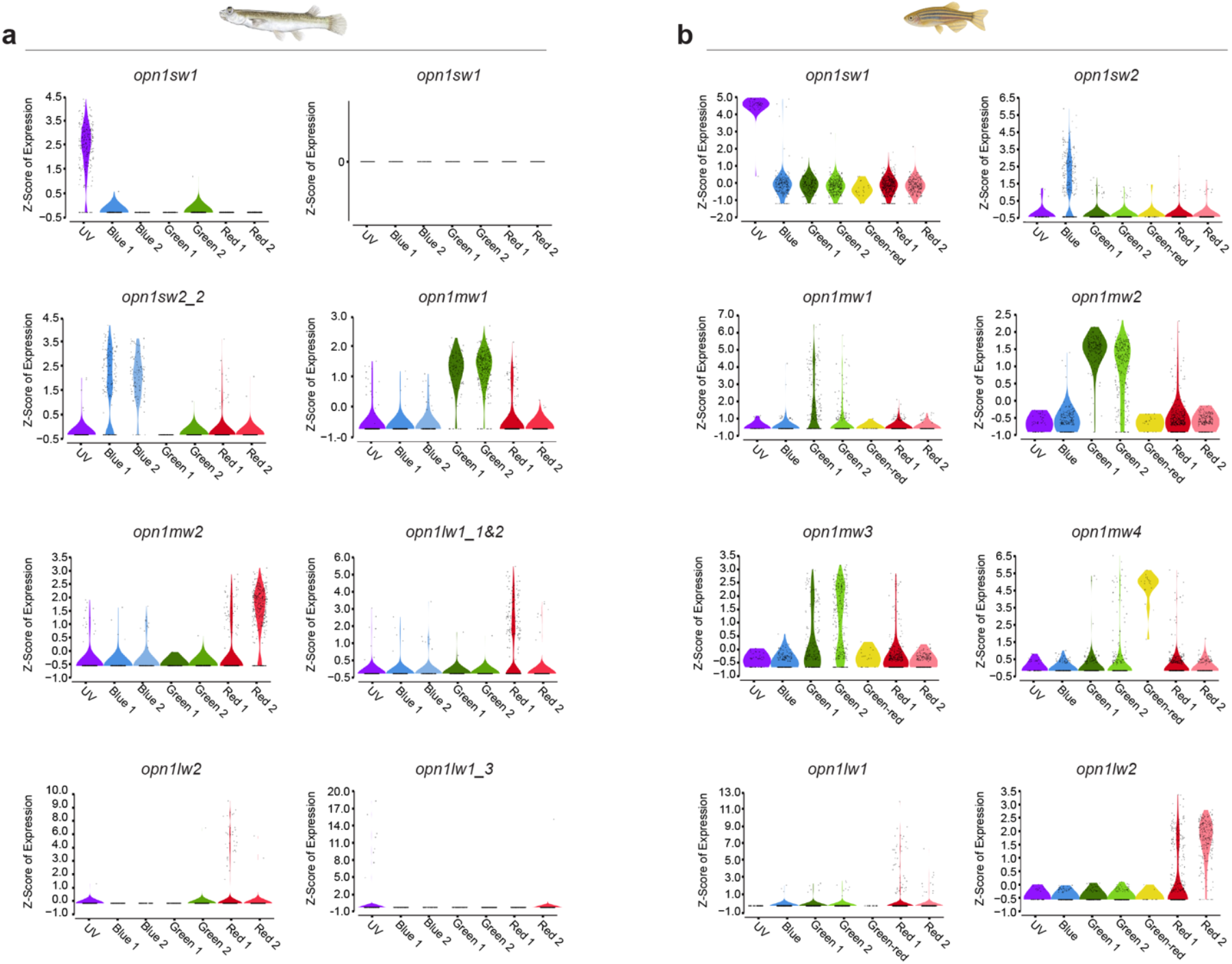
Visual opsin gene expression in cone clusters of *Anableps* and zebrafish. Violin plots show expression of visual opsin genes in the subclustered cone cells of the *Anableps* snRNA-seq (**a**) and zebrafish scRNa-seq (**b**) datasets.

**Supplementary Fig. 4.**
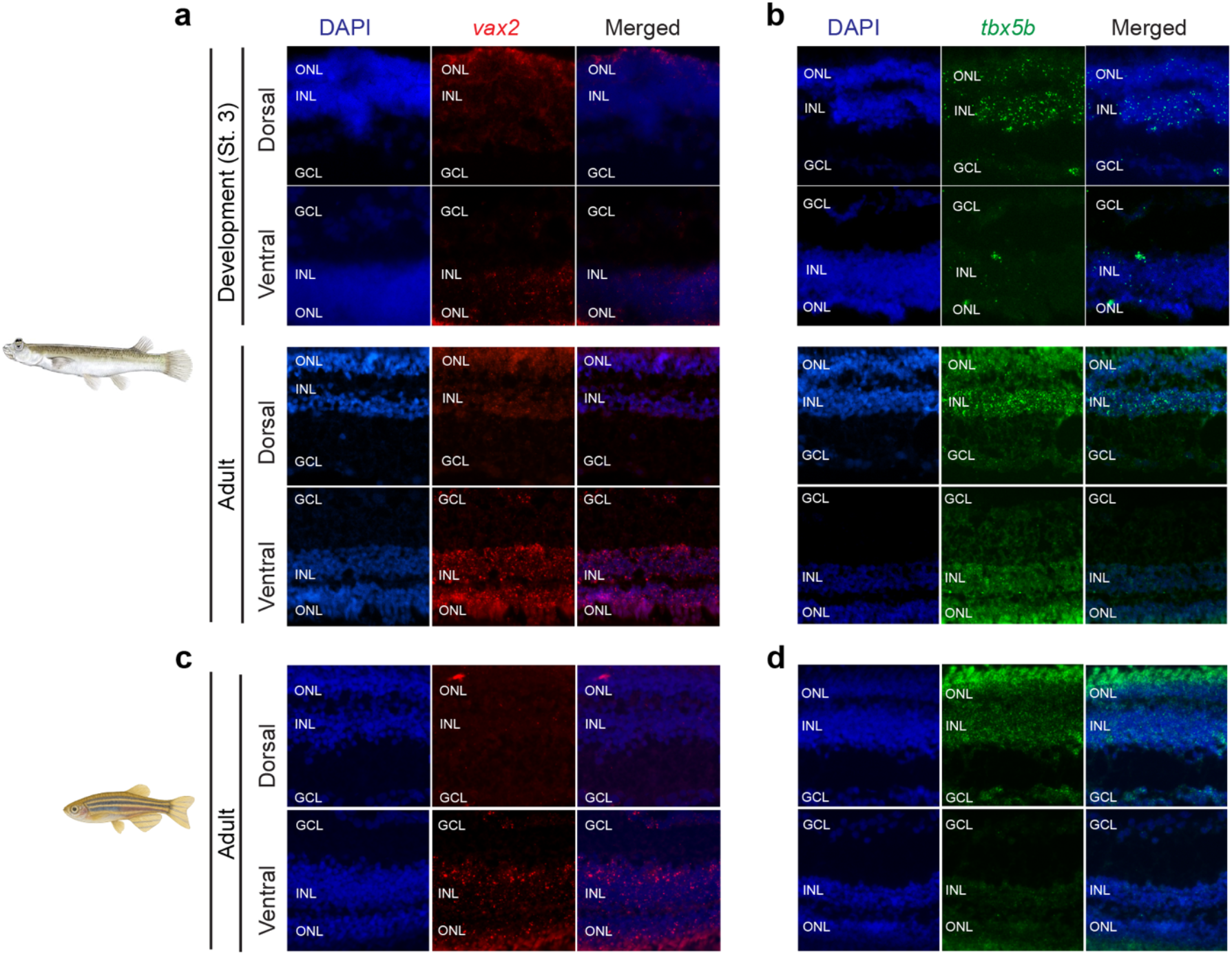
Expression of *tbx5b* and *vax2* orthologs in the dorsal and ventral retina regions of *Anableps* and zebrafish. **a-d**, HCR RNA-FISH showing expression of *vax2* (red) and *tbx5b* (green) in ventral and dorsal regions of the *Anableps* (**a, b**) and zebrafish (**c, d**) retinas. Nuclei were counterstained with DAPI (blue). ONL, outer nuclear layer; INL, inner nuclear; GCL, ganglion cell layer.

**Supplementary Fig. 5.**
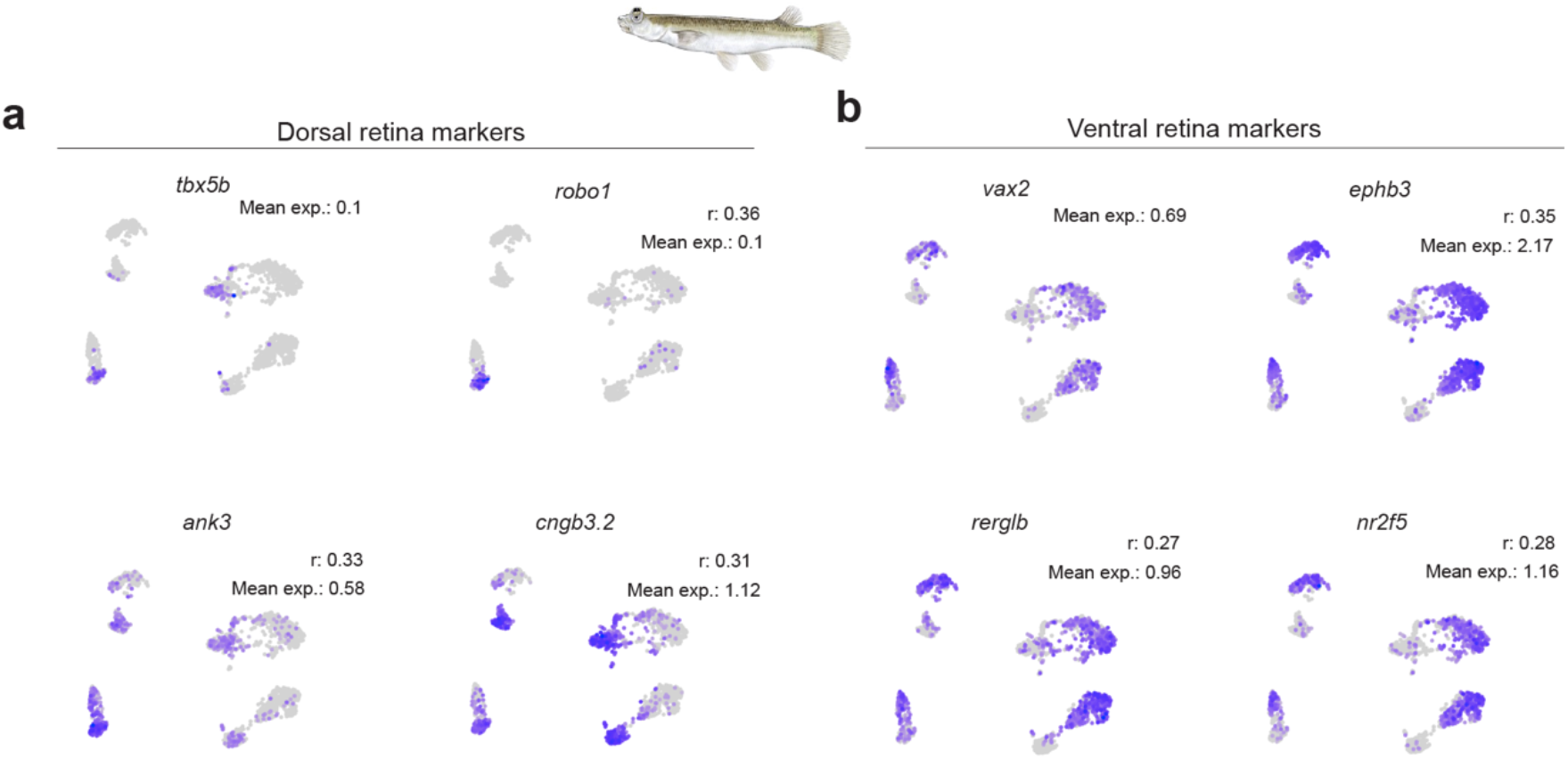
Expression of *tbx5b, vax2*, and correlated genes in *Anableps* cone cell clusters. **a**, UMAP plots showing expression of *tbx5b* and its three top correlated genes in cone cells. **b**, UMAP plots showing expression of *vax2* and its three top correlated genes in cone cells. Mean exp., mean expression; r, correlation coefficient.

## Notes

### Competing Interest Statement

The authors have declared no competing interest.

